# Estimating the conjugative transfer rate of antibiotic resistance genes: Effect of model structural errors

**DOI:** 10.1101/2021.06.29.450325

**Authors:** Sulagna Mishra, Thomas U. Berendonk, David Kneis

## Abstract

The spread of antibiotic resistance genes (ARG) occurs widely through plasmid transfer majorly facilitated via bacterial conjugation. To assess the spread of these mobile ARG, it is necessary to develop appropriate tools to estimate plasmid transfer rates under different environmental conditions. Process-based models are widely used for the estimation of plasmid transfer rate constants. Empirical studies have repeatedly highlighted the importance of subtle processes like delayed growth, the maturation of transconjugants, the physiological cost of plasmid carriage, and the dependence of conjugation on the culture’s growth stage. However, models used for estimating the transfer rates typically neglect them. We conducted virtual mating experiments to quantify the impact of these four typical structural model deficits on the estimated plasmid transfer rate constants. We found that under all conditions, the plasmid cost and the lag phase in growth must be taken into account to obtain unbiased estimates of plasmid transfer rate constants. We observed a tendency towards the underestimation of plasmid transfer rate constants when structurally deficient models were fitted to virtual mating data. This holds for all the structural deficits and mating conditions tested in our study. Our findings might explain an important component of the negative bias in model predictions known as the plasmid paradox. We also discuss other structural deficits that could lead to an overestimation of plasmid transfer rate constants and we demonstrate the impact of ill-fitted parameters on model predictions.

## 1. Introduction

Antibiotic resistance genes (ARG) and their accelerated spread in the environment have emerged as worldwide health concerns (Berendonk et al., 2015). ARG in bacteria are either located on the chromosome or on extra chromosomal elements, mainly plasmids (Bennett, 2008). Plasmids are self-replicating genetic entities that often participate in the dissemination of ARG through both vertical and horizontal gene transfer (HGT) (Zhang et al., 2011; Frost et al., 2005). Plasmids can be horizontally transmitted through conjugation, transduction, and transformation (Thomas and Nielsen, 2005), and conjugation is known to be the predominant mechanism (Norman et al., 2009). Bacterial conjugation facilitates plasmid transport from Donors (D) to Recipients (R) to form plasmid carrying Transconjugants (T) (Llosa et al., 2002). To better understand and eventually mitigate the spread of ARG, there is a necessity to quantify the rates of conjugative plasmid transfer under various environmental conditions.

The efficiency of conjugative plasmid transfer is best defined by a rate constant in a process-based model. The model can then be fitted to experimental data to obtain estimates of that rate constant. The first process-based numerical model developed by Levin et al. (1979) and its analytical solution, known as the “end-point method” (Simonsen et al., 1990), are still the most widely used techniques for estimating rate constants of plasmid transfer (Mahérault et al., 2019; Nazarian et al., 2018; Loftie-Eaton et al., 2017). There are several other studies that have developed models that can be used to estimate plasmid transfer rate constants (derivatives of the Levin et al. (1979) model, e.g., (Headd and Bradford, 2020; Kneis et al., 2019; Zhong et al., 2012, 2010), and individual-based models (interacting particle systems), e.g., (Werisch et al., 2017; Fox et al., 2008; Krone et al., 2007)). This list is not exhaustive but is sufficient to show that every conjugation model has a varying degree of complexity based on the available data, processes, and parameters included. For example, the classical mass-action ordinary differential equations (ODE) based model by Levin et al. (1979) assumed all bacterial strains grow with the same rate constant and have a bulk conjugation rate, whereas Zhong et al. (2010) took into account plasmid-bearing cost, the maturation of T, the recovery time for D after a successful conjugation event, and the formation of mating pairs. On the other hand, Headd and Bradford (2020) used a stochastic function to include the effects of the growth phase and the repression/derepression of conjugative pili synthesis in the model. Other commonly used ways to quantify transfer efficiencies are as ratios of T/D, T/R, or T/RD at a certain point in time (Wan et al., 2011; Andrup and Andersen, 1999; Richaume et al., 1989; Taylor and Levine, 1980). Since these ratios do not decouple the contributions of growth and conjugation, they are often time-dependent and sensitive to the initial experimental conditions. Therefore, though these ratios do tell us something about the transfer frequencies in an experiment, neither can they be reported as true conjugation rates nor be used for interpolation/extrapolation within the study or for comparison of transfer frequencies with other studies. Simonsen et al. (1990) also sheds light on the shortcomings of these ratios and suggests that they should not be used as indicators of plasmid transfer rates as they vary significantly based on sampling time. Similarly, the end-point solution (Simonsen et al., 1990) is also based on a single-time observation of the densities of D, R, T and, therefore, offers no provision to consider other processes (like plasmid costs or lag phases) wherever necessary.

There is currently no guideline for the factors/processes which are important to be considered when plasmid transfer rate constants are estimated from mating experiments. Therefore there is a need for a quantitative assessment of how the estimated plasmid transfer rate constants are biased if the mechanistic model used for parameter fitting has structural deficits. For example: What is the effect of ignoring plasmid-bearing costs in a conjugation model on the estimated plasmid transfer rate?

We propose virtual mating experiments (VME) to evaluate the effect of commonly encountered model structural errors on the estimate of the plasmid transfer rate constant. We used VME to study the individual and collective contribution of different additional processes towards the improvement of model fit. Virtual experiments have been used in various fields of research, with just a small number of applications in microbiology. Zurell et al. (2010) reviewed various studies that have used the virtual experiments approach in different fields of ecology. The authors described virtual experiments as a powerful approach to optimize sampling designs and modeling tools. Earlier Simonsen et al. (1990) used computer simulations to study the effects of factors like D:R ratios, lag phases, growth rate differences on the efficiencies of their proposed end-point method and reported the circumstances under which the method might be unsuitable for use. However, this sensitivity analysis was conducted only for the first three hours of the exponential phase; hence the finding might not be transferable to the entire duration for longer experiments.

To our knowledge, this is the first study that quantifies the effect of model structural deficits on the estimates of plasmid transfer rate constants in liquid culture by means of VME. Even though most of the ARG found in the environment are likely to reside in biofilms, we designed our study based on conjugation experiments for well-mixed liquid culture. This is because mating experiments in liquid culture are still widely used in research and convenient for analyzing the effect of various environmental/experimental factors on the behavior of populations (e.g., multi-well conjugation experiments). We study the impact of the following four factors: (a) the plasmid cost, (b) the lag phase in growth, (c) the dependency of plasmid transfer rates on population density, and (d) the maturation time of newly formed T strain. For running the VME, we used a forward conjugation model taking into account all the factors a-d and collected virtual samples from the model output at different temporal resolutions. We then fitted “inverse” models with different structural deficits (e.g., considering only processes a and b while neglecting c and d) to the virtual samples to obtain an estimate of the plasmid transfer rate constant. We finally compared the estimated transfer rate constants to their known “true” values so as to assess the impact of the model’s structural deficits. We verified the outcomes under variable experimental conditions and sampling strategies typically encountered in microbial practice.

We preferred a VME based study over an empirical one in order to suppress inevitable experimental noises. This helps to explicitly study the effect of model structural errors on the estimated plasmid transfer rate constant. Since the “true” values of all the variables, process rates, and parameters are known, the VME approach allows us to make bold inferences about the efficiencies of the different inverse models and sampling schemes. Furthermore, the configuration of both forward and inverse models can be easily altered to analyze a wide range of scenarios.

Specific objectives of the study were: a) To quantify the bias in estimated rate constants of plasmid transfer arising from structural model errors, b) To analyze the sensitivity of the estimated parameter (obtained from different structurally deficient models) to experimental conditions (covering real-world scenarios), and c) To demonstrate how VME can help with optimization of sampling schemes.

## 2. Methods

### 2.1. VME approach

The procedure is divided into three major steps: 1. the forward model (FM), which serves as a surrogate for the real microbial system, 2. the virtual sampler, which collects data under given sampling frequencies, 3. the inverse model (IM), which is fitted to the virtual data to obtain an estimate of the plasmid transfer rate constant.

The effect of structural model deficits is represented by the bias in the estimate of the rate constant where the “true” value (as used in the forward model) is known. The three steps are then repeated for variable experimental conditions and sampling frequencies (Fig. 1). To avoid confusion, the densities of the bacterial strains generated by the FM will be referred to as “virtual observations”.

**Figure 1:**
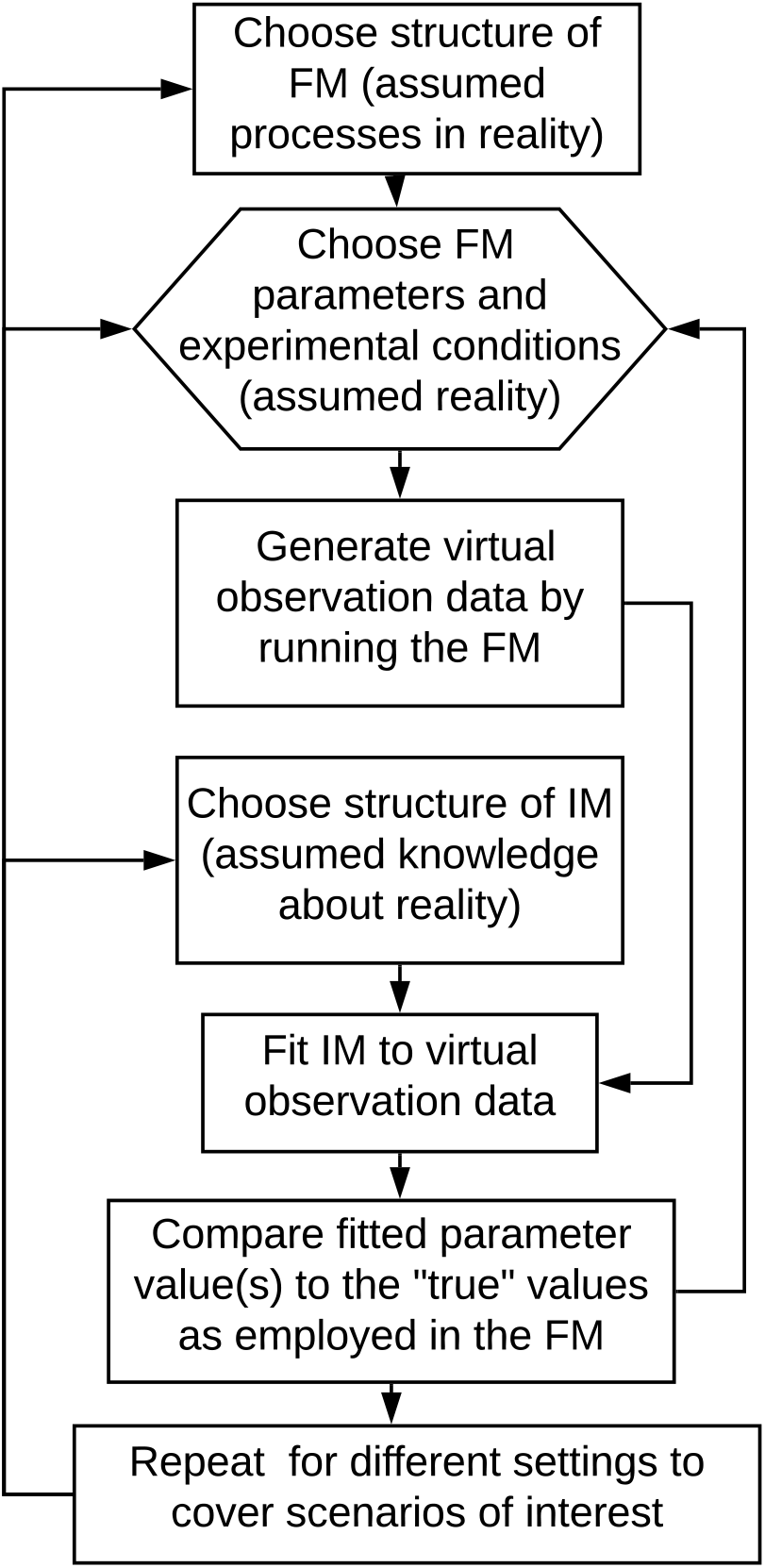
Flow chart summarizing the steps followed in the VSE study. The initial concentrations of the four state variables and six parameters (known from previous studies) are used as inputs for the Forward model (FM). Samplings at pre-determined frequencies are conducted from the generated “truth”. The inverse model (IM) is fitted to this data for parameter estimation under ten modeling scenarios and the results are compared with the known or “true” parameter values. The overall results are validated for different experimental conditions.

### 2.2. Model configurations

#### 2.2.1. Forward model

The core of the forward model is given by the set of differential equations (Eq. 1) proposed by Levin et al. (1979).

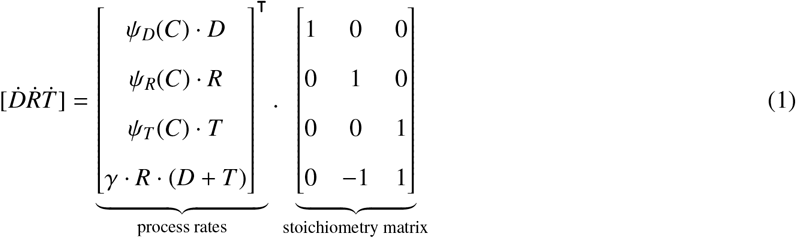

In Eq. 1, the left-hand side vector contains the derivatives of the cell densities of the individual strains (D, R, and T) with respect to time (cells ml^−1^ hour^−1^). The process rates vector accounts for the resource-limited growth rate of the strains (C denotes the resource) *ψ_strain_* in rows 1-3 and a bulk plasmid transfer rate constant *γ* (ml cells^−1^ hour^−1^). In most cases, resource limitation is expressed by a Monod model or a carrying capacity. For simplicity, Levin et al. (1979) model assumed all strains share the same intrinsic growth rate constant (i.e., *ψ_D_*=*ψ_R_*=*ψ_T_*=*ψ*). The FM in our study is developed by extending Eq. 1. We added four processes to the original model and investigated the contribution of each one of them towards the accuracy in the estimated plasmid transfer rate constant.

- Process 1 - (*CST*: takes into account the reduction in the growth rate due to plasmid-bearing costs (Carroll and Wong, 2018).
- Process 2 - (*LAG*): caters to the initial “no-growth” phase encountered when the D and R are confronted with new experimental conditions. This is otherwise known as the initial adjustment phase or inactive stage, or the lag phase in growth (Rolfe et al., 2012). We assume that the cells newly formed during the course of our experiment do not have to undergo an additional no-growth phase as there is no change in the experimental conditions and medium.
- Process 3 - (*DNS*): introduces a dependency of plasmid transfer on growth. Therefore, we assume a decline in the plasmid transfer rate as the system approaches the stationary phase, e.g., due to resource limitation, as reported by empirical studies (Levin et al., 1979; Sysoeva et al., 2020).
- Process 4 - (*MAT*): includes the maturation time required for a newly formed T to acquire the full capabilities of a donor cell. Immature T cannot transmit the plasmid to further recipients (Zhong et al., 2010).

The FM, including all the four processes listed above, is expressed in Eq. 2. In our model, the processes are represented in the form of functions with the same names as the processes (*CST, LAG, DNS, MAT*) and are explained in Eq. 3.

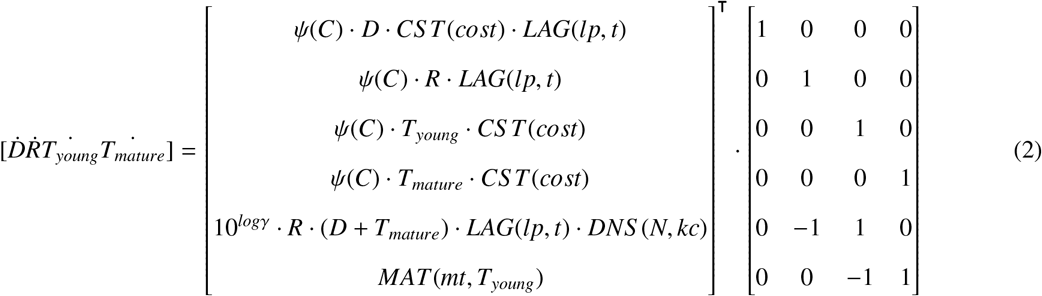

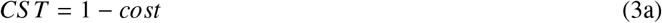

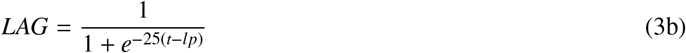

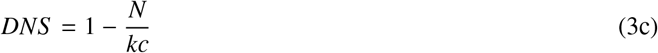

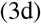

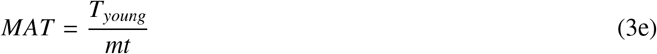

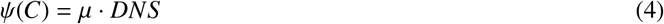

where *t* stands for time (hours), *N* stands for the total density of the population (D+R+T), total transconjugants T = T*_young_*+T*_mature_*. All the parameters occurring in Eq. 3 and 4 are defined in Table 1. Eq. 3b is based on the mechanistic model for simultaneously determining the lag phase (Huang, 2011, 2008). Knowing that conjugation and growth stage are highly co-related (Schuurmans et al., 2014), we used the same *LAG* for the plasmid transfer process in our model (Eq. 2). Note that the newly introduced state variable T*mature* represents the density of the mature transconjugants that are capable of infecting further recipients. T*young* used in Eq. 3e in function *MAT* represents immature or newly formed T. The resource-limited growth of the strains is implemented using the carrying capacity of the medium (Eq. 4). The initial densities of D, R, *T_young_, T_mature_* were set to 10^4^, 10^4^ 0, and 0 CFU ml^−1^. We ran our virtual mating experiments for 12 hours. Note that for the proper scaling of the parameters, we used *γ* in log_10_ units. We chose parameter values representative for *E. coli* because of its relevance in the context of the spread of ARG in different mediums (Huddleston, 2014; Teuber, 1999) and a good amount of prior knowledge (Kneis et al., 2019; Nazarian et al., 2018; Zhong et al., 2010; Simonsen et al., 1990). Nevertheless, based on the objective, species, and plasmid under study, the processes/ parameters can be altered or adapted while the overall VME framework stays intact.

**Table 1:**
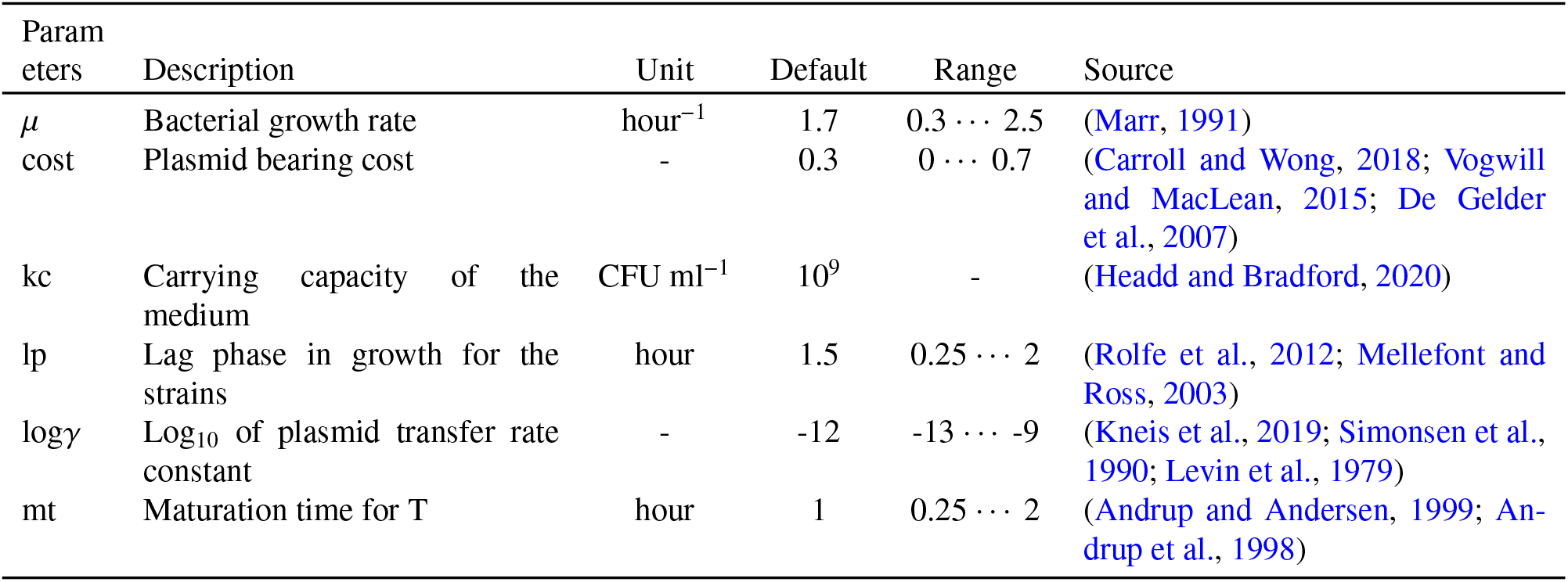
Parameter default values (used for evaluation of the modeling scenarios and sampling frequency) and their ranges (used for validation of the results)

#### 2.2.2. Numerical solution of the ODE

We implemented the FM and the IM in R (Team, 2011). For computational convenience, we used the R package - *rodeo* (Kneis et al., 2017) to implement the ODE-based models. For numerical integration, *rodeo* uses the package *deSolve* (Soetaert et al., 2010).

#### 2.2.3. Inverse model

##### Modeling scenarios

To find the effect of various model deficiencies on the error in estimated plasmid transfer rate constants, we systematically increased the model complexity from the basic Levin’s model (Eq. 1) to the forward model (Eq. 2). For example: When FM is scenario 10 (that includes all four processes) and IM is scenario 6 (that includes only LAG and CST); biases in the estimated log*γ* inform us about the impact of structural deficits (DNS and MAT) on the estimate of transfer rate constant.

##### Sampling

The virtual sampler conducted sampling according to two pre-defined strategies: (a) Fixed frequency of sampling where we collected samples every 15 minutes to every 6 hours, and (b) Fixed number of samples per bacterial doubling time where we varied the number of samples between 1 and 4 per doubling time (e.g., at *μ* = 0.7 hour^−1^, doubling time ≊ 1 hour, samples are collected at the rate of 1 - 4 samples/hour). The virtual sampler collects the densities of D, R, and total T because T*young* and T*mature* cannot be distinguished in practice.

##### Parameter estimation

Using inverse modeling, the different IMs (Tab. 2) were fitted to the observed time-dependent cell counts of D, R, T (virtual observations) to estimate the plasmid transfer rate constant (log*γ*). The sum of squared residuals was chosen as the objective function. Cell densities were log-transformed prior to the calculation of residuals. Observations of D, R, and T were equally weighted.

**Table 2:** List of additional processes included in the modeling scenarios (IM). Refer Eq. 3 and Sec. 2.2.1 for the function definitions.

| Scenarios | Additional process included |
| --- | --- |
| 1 | None (identical to Eq.1) |
| 2 | *CST* |
| 3 | *LAG* |
| 4 | *DNS* |
| 5 | *MAT* |
| 6 | *CST*, *LAG* |
| 7 | *CST*, *DNS* |
| 8 | *CST*, *LAG*, *DNS* |
| 9 | *CST*, *DNS*, *MAT* |
| 10 | All (identical to the FM) |

### 2.3. Evaluation and validation

#### Scenario evaluation

We evaluated the performance of the different models based on the error in the estimated plasmid transfer rate constant with respect to its known value used in the FM (log*γ_estimated_* - log*γ_true_*). A comparison of the errors in the parameter estimates under different sampling frequencies was made to identify a suitable sampling protocol. Inspection of the residual model errors helped us to identify the most suitable model for estimating the log*γ* in well-mixed cultures under typical experimental conditions (Tab. 1).

#### Sensitivity analysis

We verified the transferability of results from the previous section for variable experimental conditions. We carried out a sensitivity analysis by varying the parameters assumed in the FM: *μ*, log*γ*, lp, mt, and cost (Tab. 1) one at a time while the other parameters were fixed at their default values (also Tab. 1). We computed the error in the estimated log*γ* with respect to each of these parameters for a fixed sampling frequency. We conducted the sensitivity analysis to cover most real-world scenarios and defined the parameter ranges based on the values observed in external empirical studies (See Tab. 1). We also studied the effect of the initial D:R ratio on the error in the estimated log*γ* by varying D:R between 1000:1 to 1:1000 in factors of 10. We analyzed our results in two stages: a) the contribution of each function/process towards reducing the error in estimated plasmid transfer rate (scenarios 2 - 5) and b) their dynamics when added to a model in different combinations (scenarios 6 - 9). We ran each test case for all the modeling scenarios (10 IMs), where scenarios 1 and 10 were used as controls (basic and best-case scenarios).

## 3. Results

### 3.1. Modeling Scenarios

#### 3.1.1. Effect of sampling frequencies

The frequency of sampling (varied between 15 minutes and 3 hours) had a minor impact on the accuracy of the estimated log*γ* for all the modeling scenarios, which include *LAG* (Fig 2). However, other scenarios 1, 2, 4, 5, 7, and 9 showed an improvement in parameter estimation as the number of samples collected decreased. While more data usually results in better fits, we observed the opposite effect in scenarios with missing *LAG*. This was because, with frequent sampling from the lag phase, the fitting of the inadequate models (with a missing *LAG*) resulted in a much poorer fit. Scenarios 1 - 5 consistently underestimated the log*γ*. At a sampling frequency of 2 hours, exclusion of all four processes (*CST, LAG, DNS, MAT*) resulted in an underestimation of the plasmid transfer rate constant by 2.3 orders of magnitude. While individually LAG, *CST*, and *DNS* reduced the error in the estimated parameter by 1.3, 0.9, 0.4 order of magnitude, *MAT* did not show any notable impact (Fig. 2a). With the default test settings (Tab. 1), the errors in estimated *γ* were the highest (by 3.2 orders of magnitude for sampling done every 15 minutes) in scenario 1, which was the basic model from Eq. 1 without any additional processes.

**Figure 2:**
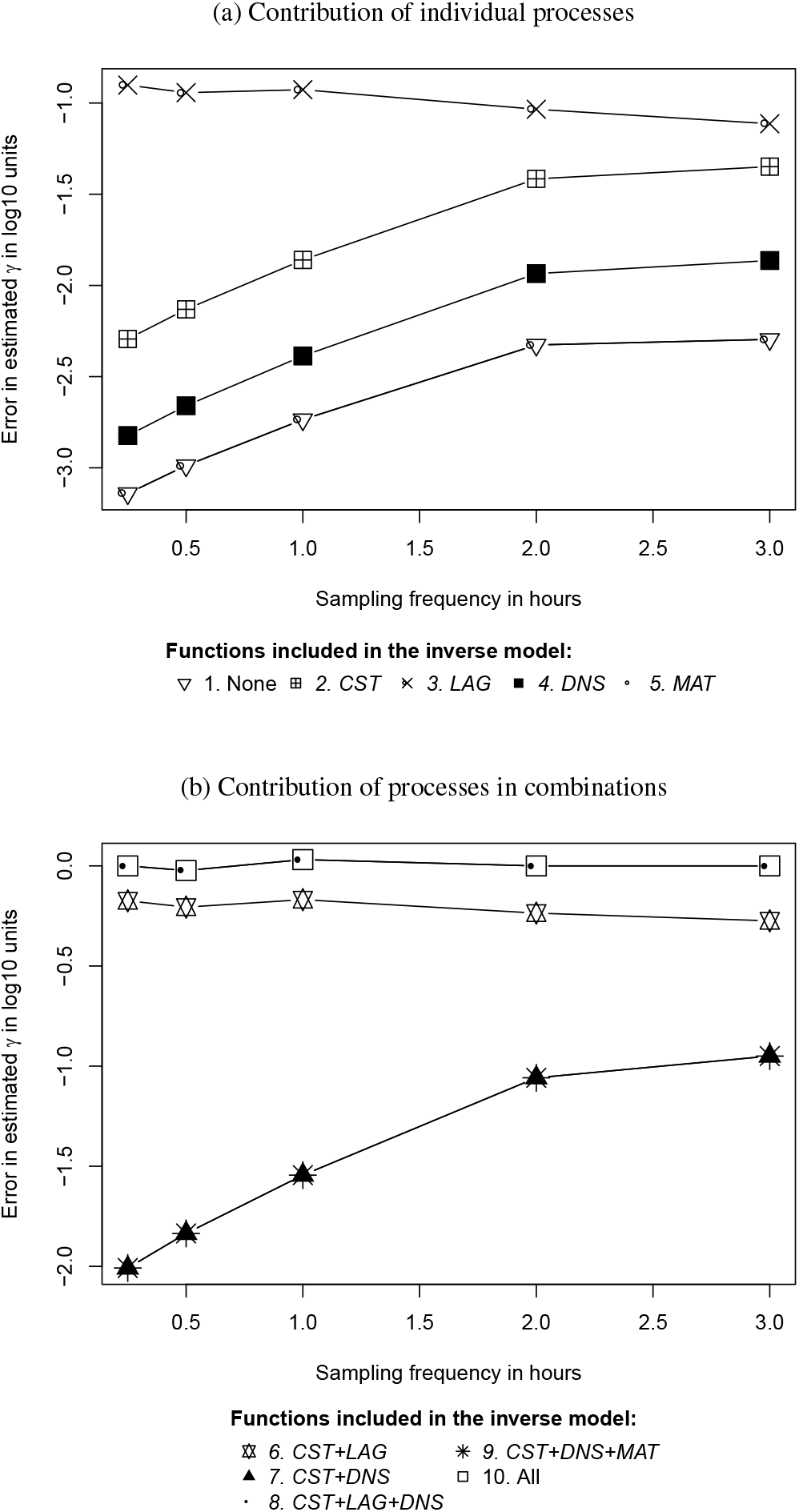
Error in estimated plasmid transfer rate constants (log*γ*) for each modeling scenario at different sampling frequencies. All results refer to the forward model (scenario 10) ran with default settings. The legend lists scenario number and additional functions included in each one over the Levin et al. (1979) model. Function definitions- *CST*: Plasmid cost, LAG: Lag phase, *DNS*: Density dependency *MAT*: Maturation time. Refer to Sec. 2.2.1 for details of the function definitions. Note the difference of scale in the Y-axis. Negative errors indicate an underestimation of the log*γ* by structurally deficient models.

The errors in the estimated log*γ* in scenarios with combinations of processes (scenarios 6 - 10 Fig. 2b) were in agreement with the results from the individual processes (Fig. 2a). At the sampling frequency of 15 minutes, scenarios 7 and 9 recorded errors of ≈ 2 orders of magnitude of estimated *γ* while 6 and 8 resulted in negligible errors. *MAT* and *DNS* continued to have little impact on the estimates also when used in combination. Among all structurally inadequate IMs, scenario 8 was the best performing model (error ≈ 0), closely followed by scenario 6, which resulted in an estimation error of 0.2 orders of magnitude.

We computed the error in the estimated log*γ* for all scenarios with a gradually decreasing sampling frequency of up to only 1 sample every 8 hours. For samples collected in intervals greater than 3.5 hours (not included in the figure), insufficient virtual observations led to the non-identifiability of the log*γ*. We verified the results for different assumed bacterial growth rates. Keeping the other parameters fixed to the default settings (Tab. 1), we found that for doubling times of 20 minutes to 4 hours, it is sufficient to take samples every 2 hours for estimating the log*γ* in liquid culture experiments (Fig. 2a). The alternate sampling arrangement based on the growth rate of the strains (See section 2.2.3) showed that it is sufficient to collect 1 sample every 4 - doubling time (for cultures with *μ_R_* = 1.7 hour^−1^) in order to obtain an unbiased model fit for estimating the log*γ*.

#### 3.1.2. Comparison of virtual observations with simulations

The strain densities calculated using the IMs are referred to as “simulations”, which we used to highlight the effect of using structural deficit models for the prediction of the D, R, and T populations. The performance of the different model structures (Tab. 2) in estimating the densities of the three bacterial strains varied from one another (Fig 3). The virtual observations were compared with the simulations generated from scenarios 1 - 9 for two plasmid transfer rate constants (a) log*γ*= −12 (default value) and (b) log*γ*= −9. The other parameters were fixed to the default values from Table 1 and samples were collected every 2 hours. The densities of observed and simulated D and R were similar in both cases because their densities are predominately governed by growth and the conversion of R into T has little effect on the total number of R cells. These plots highlight the importance of using structurally correct models to get unbiased estimates of the population densities.

**Figure 3:**
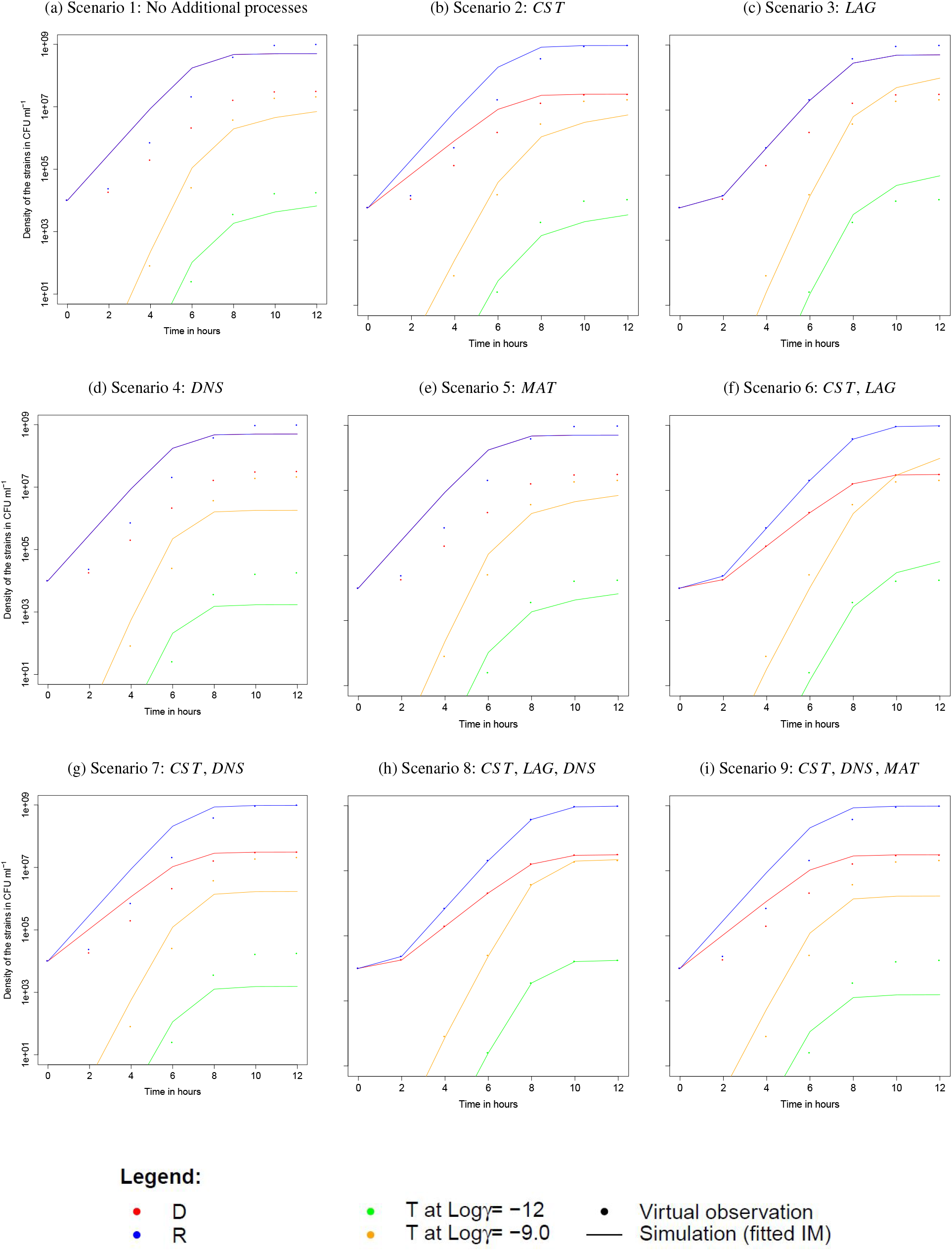
Comparison of virtual observations to simulations for the strains D, R, and T for each modeling scenario for virtual sampling done every 2 hours. Note that T (in green and orange) are shown for *γ* = −12 and −9 in log_10_ units. The observations (from the FM) and simulations (from the IM) for D and R for both values of log*γ* are almost identical and cannot be visually distinguished. Results for scenario 10 have been omitted as it gave a perfect fit (IM identical to FM). Function definitions- *CST*: Plasmid cost, LAG: Lag phase, *DNS*: Saturation *MAT*: Maturation time.

Scenario 1-5 overestimated the densities of D, R, T in the exponential growth phases, the exception being scenario 3, where the estimated D and T showed no notable bias. In both cases (a and b), scenario 8 (which included the plasmid cost, the lag phase, and the dependency of plasmid transfer on population density) produced the best fit results with respect to all three strains. Scenario 6 (which included the plasmid cost and the lag phase) provided good estimates for R and D for both cases, while the T was slightly underestimated in the exponential phase. For the rest of the scenarios, R and D are consistently overestimated by the models until the stationary phase is reached, except for the anomaly observed in scenario 1 (no additional processes) - where the D strain is grossly overestimated throughout the simulation (not visible because the graph collapses with that of the R strain). In all scenarios (except 3, 6, and 8), T was overestimated until saturation and underestimated afterward. We did not find a notable effect of the value of log*γ_true_* (assumed in the FM) on the performance of the models in estimating the densities of the strains.

### 3.2. Sensitivity analysis

#### 3.2.1. Contribution of the individual processes

##### Effect of the plasmid cost

The error in the estimated *γ* obtained for scenarios 1, 3, 4, and 5 (where *CST* is excluded) increased by over 2 orders of magnitude when the plasmid-bearing cost increased from 0 to 0.7. The error in scenario 2 (including *CST*) reduced by 0.5 orders with increasing plasmid cost, thus highlighting the importance of including the plasmid cost in a conjugation model (Fig. 4a). However, at low plasmid costs (< 0.2), scenario 3 (including *LAG*) and scenario 4 (including *DNS*) outperformed all other IM scenarios. We analyzed the performance of an IM in terms of the improvement in the estimated log*γ* (due to the inclusion of one or more processes) with respect to scenario 1 (with no additional processes, Eq. 1).

**Figure 4:**
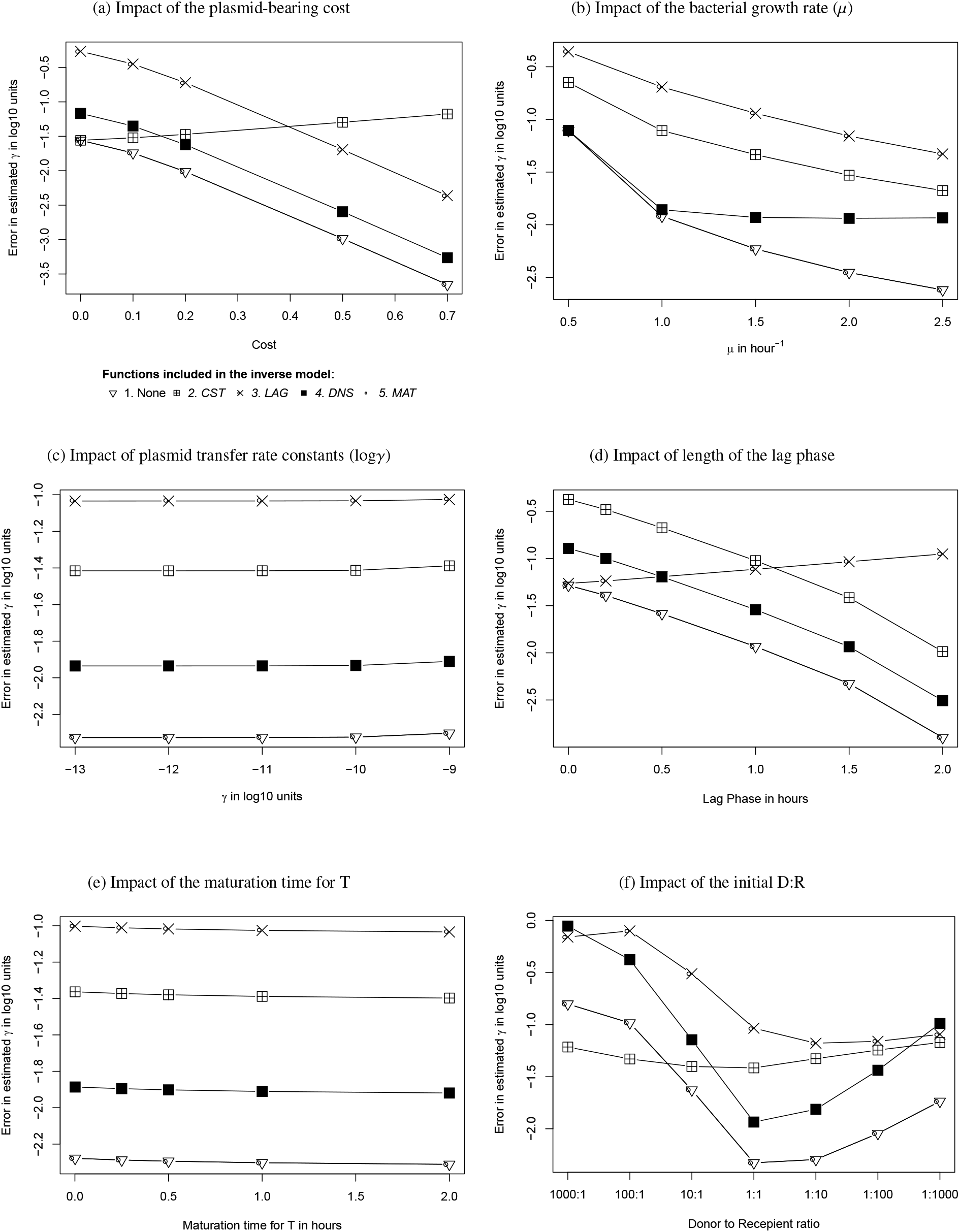
Sensitivity of model outcomes (in terms of the error in estimated log*γ*) to variable initial parameters for scenarios 1 to 5 (with single processes). Samples from the FM were collected every 2 hours. Note that for each study (subfigures a-e), only the one mentioned parameter was altered from the default settings. In subfigure (f), Sensitivity analysis was performed with respect to a variable initial D:R ratio while the parameters were the same as for the default settings (Tab. 1). Negative errors indicate an underestimation of the log*γ* by structurally deficient models. Function definitions- *CST*: Plasmid cost, *LAG*: Lag phase, *DNS*: Saturation *MAT*: Maturation time.

##### Effect of μ

The performance of all modeling scenarios (1 - 5) decreased as the growth rate constant *μ* increased from 0.5 to 2.5 hour^−1^ (Fig. 4b). At lower growth rate constants (< 1 hour^−1^), including the dependency of conjugation on population density (*DNS*), showed no pronounced impact on the model performance (compare scenarios 1 and 4 in Fig. 4b). At higher growth rates (*μ* ≥ 2.3 hour^−1^), the inclusion of function *DNS* improved the estimates by ≈ 0.7 order in comparison to 0.4 orders at *μ* = 1.7 hour^−1^ (default settings). The maximum error in estimated log*γ* observed at low growth rates (*μ* = 0.5 hour^−1^) was 1 order of magnitude.

##### Effect of the logγ

For the tested range of log*γ* values, the output of the different modeling scenarios was essentially insensitive to varying log*γ* (Fig. 4c). The role of each function in reducing the estimation error followed the order *LAG*> *CST*» *DNS* > *MAT* contributing ≈ 1.3, 0.9, 0.4, and 0 each.

##### Effect of the lag phase

As expected, we observed an increased relevance of *LAG* in the models as the length of the lag phase in the FM increased from 0 to 2 hours (Fig. 4d). The pattern remained consistent until the lag phase of 4 hours (not included in the results). Under default settings, the maturation time (*MAT*) was not important to be included for any length of the lag phase. At shorter lag phases (< 0.5 hours), scenario 2 (including *CST* only) and 4 (*DNS* only) outperformed scenario 3 (*LAG* only).

##### Effect of the maturation time

At the default value of log*γ* = −12, the change in the maturation time showed no effect on the estimated log*γ* (Fig. 4e). The error in estimation increased with the maturation time in all models which were subjected to structural deficits (scenarios 1-5). Under default settings, the contribution of the single functions in reducing the estimation error followed the order *LAG* > *CST* » *DNS* (> *MAT*≈ 0).

##### Effect of the D:R ratio

While the errors in the estimated parameter in scenario 2 (including *CST*) remained fairly constant with the variation in the initial D:R ratios, scenario 3 (including LAG) showed a variation of < 1 order in the estimated error between the extremes (compare D:R 1000:1 and 1:1000 for scenario 3 in Fig. 4f). Other scenarios recorded high variability in the estimation errors ≈ 2 orders between the different test conditions (Fig. 4f). For scenarios 1, 4, and 5 the error in estimation followed the D:R order 1:1 > 1:1000 > 1000:1. At high D:R ratios (> 100:1), scenario 2 was the worst-performing model. For the rest of the conditions (D:R < 10:1), the contribution of the processes in reducing the error in estimated log*γ* followed the order *LAG* > *CST* » *DNS* (> *MAT≈* 0).

#### 3.2.2. Contribution of processes (in combinations)

##### Effect of the plasmid cost

At all plasmid-bearing costs, scenarios 6, 8 (and 10) recorded the minimum error in estimated *γ* (Fig. 5a). *MAT* had a negligible contribution towards the improvement of the model performance. The estimation of log*γ* obtained for scenarios 7 and 9 (where *LAG* is excluded) improved by 0.4 orders as the assumed plasmid-bearing cost increased from 10 - 70%. Scenario 6 (CST+LAG) reduced the error in estimated *γ* by 2.4 orders of magnitude when the assumed plasmid cost was 0.7 in comparison to 1.4 orders when the cost was 0.1.

**Figure 5:**
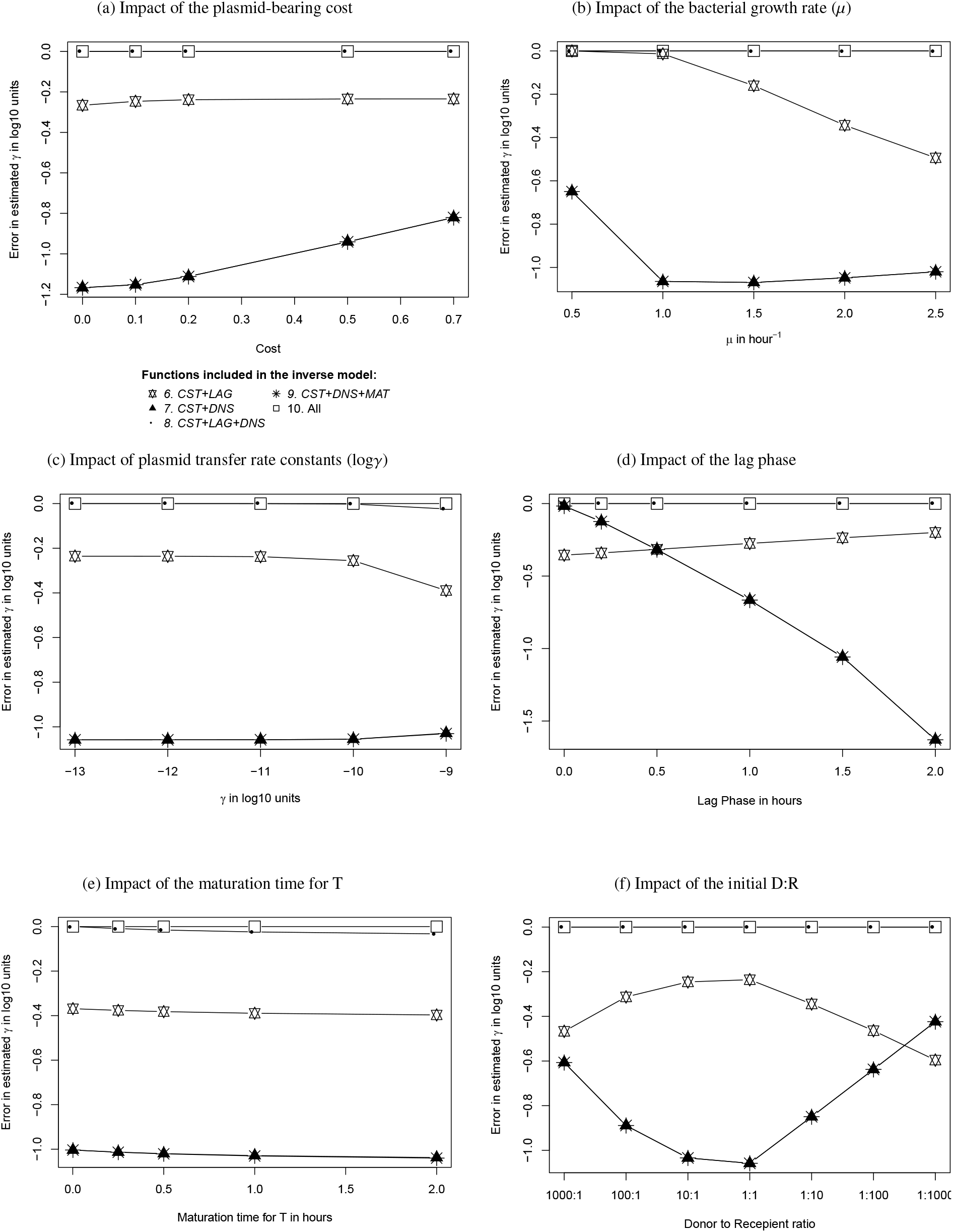
Sensitivity of model outcome (in terms of the error in estimated log*γ*) to variable initial parameters for scenarios 6 to 10 (with combinations of processes). Samples from the FM were collected every 2 hours. Note that for each study (subfigures: a-e), only the one mentioned parameter was altered from the default settings. In subfigure (f), Sensitivity analysis was performed with respect to a variable initial D:R ratio while the parameters were the same as for the default settings (Tab. 1). Negative errors indicate an underestimation of the log*γ* by structurally deficient models. Function definitions- *CST*: Plasmid cost, LAG: Lag phase, *DNS*: Saturation *MAT*: Maturation time.

##### Effect of μ

At low growth rate constants (< 1.5 hour^−1^), scenario 6 showed negligible error in the estimated parameter and a maximum error of 0.4 orders for extremely fast-growing cultures with doubling time < 20 minutes (Fig. 5b). At lower growth rate constants (< 1.0 hour^−1^), including the dependency of conjugation on population density (*DNS*), had no pronounced impact on the model performance (compare scenarios 6 and 8 in Fig. 5b). At higher growth rates, excluding *LAG* (scenarios 7 and 9), resulted in an increase in the error in the estimated log*γ*.

##### Effect of the logγ

At lower plasmid transfer rates (e.g., log*γ* < −10), the combination scenarios were insensitive to varying log*γ* (like the individual functions in Fig. 4c). However, as conjugation became a relevant process (log*γ* > −10), we observed a rise in the contribution of *DNS* (compare scenario 6 with 8 in Fig. 5c).

##### Effect of the lag phase

Scenarios 7 and 9 (where *LAG* is omitted) recorded a steep increase in the estimation error (from 0 to 1.7 orders of magnitude) with the duration of the lag phase (Fig. 5c). At shorter lag phases (< 0.5 hours) scenario 7 (*CST, DNS*) outperformed scenario 6 (*CST, LAG*).

##### Effect of the maturation time

For all tested values, the maturation time had no impact on the performance of the various IMs (Fig. 5e). Including *DNS* in scenario 6 (resulting in scenario 8) improved the estimate by 0.4 orders, whereas including *LAG* to scenario 7 (also resulting in scenario 8) improved the estimate by ≈ 1 order of magnitude.

##### Effect of the D:R ratio

While the errors in the estimated parameter in scenarios 8 (and 10) remained constant with the variation in the initial D:R ratios, scenarios 6 showed a negligible variation of < 0.2 orders between the extremes (Fig. 5f). However, estimates from scenarios 7 and 9 varied over > 0.6 orders of magnitude with initial D:R ratios.

## 4. Discussion

Ignoring any of the four processes *CST, LAG, DNS* or *MAT* generally leads to an underestimation of the log*γ* in the modeling scenarios. The importance of the processes generally followed the order *LAG* > *CST* > *DNS*, whereas; *MAT* only had a negligible impact on the estimates in the range of environmental conditions tested in this study. Below we discuss the individual and combined effect of these processes and how their relative importance varies with experimental conditions.

### Cost (CST)

A reduction in intrinsic growth rate constants due to plasmid carriage is a typical phenomenon. Several studies have reported up to > 50% reduction of growth in plasmid-bearing cells compared to plasmid-free recipients (He et al., 2019; Carroll and Wong, 2018; Vogwill and MacLean, 2015). If the inverse models ignore plasmid costs, the model assumes all strains to grow at the same unimpeded rate (See, e.g., Eq. 1). When such a model is fit to the virtual observations, the overestimation of the growth rate of the D and T strains is compensated by a negative bias in the plasmid transfer rate constant. Hence in modeling scenarios 1, 3, 4, and 5, we observe that the densities of strain D are overestimated with respect to the observations (Fig. 3). A lower log*γ* further lead to the overestimation of R densities.

Under default settings (Tab. 1), adding *CST* to the model reduced the error in the parameter estimate by ≈ 3 orders of magnitude at a high plasmid cost (> 0.5). The difference between the true and the estimated *γ* improved by > 0.7 orders of magnitude when *CST* was taken into account, even when the plasmid cost was assumed to reduce the growth of D and T by only 20%. Therefore, we consider the inclusion of the plasmid cost to be inevitable for the proper inference of plasmid transfer rate constants.

It is known from previous studies that initial D:R ratios should not have an impact on the estimated plasmid transfer rate constants (Simonsen et al., 1990). *CST* played the most important role in estimating the population dynamics, mainly in conditions with D:R > 100, where the growth of D and T are grossly overestimated when the plasmid cost was not taken into consideration. The log*γ* estimated using the scenarios with *CST* were less sensitive to initial D:R. In the other scenarios, we observed a large dependence of the estimated log*γ* on initial D:R ratios.

Though we observed an underestimation in the log*γ* due to the neglect of *CST*, we are aware of some studies that reported a growth advantage for some plasmid-bearing cells (Wu et al., 2018). For plasmids that have a positive impact on the growth of D and T, we would observe an overestimation in the plasmid transfer rate constant when *CST* is neglected. In such a situation, the overestimation caused by neglecting *CST* can be countered by the exclusion of *LAG* (which always creates a negative bias) to produce an IM which could give unbiased estimates of the transfer rates, despite the structural deficits. However, VME must be used to quantify the bias in the parameter estimates arising from such structural deficits (as an individual deficit or in combination) to ensure an apprised model development.

### Lag phase (LAG)

Studies suggest that the lag phase in bacteria could last from anything between 20 minutes to a few hours (Rolfe et al., 2012; Mellefont and Ross, 2003). In the absence of LAG, the growth parameter (*μ*) is wrongly assumed to be constant throughout the simulation. This leads to an overestimation of the densities of R and D. The inevitable overestimation of T, however, is prevented by an underestimation of the log*γ* in the fitting process. Under the default parameterization of FM, *LAG* was found to reduce the total error in the parameter estimate by 1.3 orders of magnitude (which is ≈ 60% of the total error). In systems with a shorter lag phase (e.g., < 30 minutes), the error in the estimated log*γ* due to the omission of *LAG* is of minor importance only (bias in the estimated parameter < 0.5).

### Saturation (DNS)

Empirical studies have reported that the log*γ* reduces by ≈ 2 orders of magnitude as the mating culture approaches the stationary growth phase (Levin et al., 1979). Function *DNS* defines the conjugation process rate to be dependent on resource availability (or, in our case, total cell density. Eq. 3c). Ignoring *DNS* allows for constant high production of T throughout the simulation, which is necessarily compensated by a negatively biased estimate of the log*γ*. Under the default parametrization, the inclusion of *DNS* reduced the total error in the estimated log*γ* by only 0.4 orders of magnitude. At lower growth rates < 1.5 hour^−1^, *DNS* was found to be of minor importance because in such cultures, the stationary phase is reached much later and the problem of an overestimation of T is no longer important unless the mating experiments continue for more than 12 hours. On the other hand, when the growth rate or the initial total cell density was high, the relevance of including *DNS* in the IM also increased because a larger part of the observations now fell in the stationary phase.

### Maturation time (MAT)

*MAT* introduces a delay in the number of T capable of plasmid transfer, which results in a reduction of conjugation activity. When *MAT* is ignored in an inverse model, the entire pool T (young and mature) is assumed to participate in plasmid transfer, which necessarily causes a negative bias in the estimated plasmid transfer rate constant. However, for the maturation time of T chosen between 0 - 2 hours, including *MAT* in the model had no effect on the estimated log*γ*. This is because, under typical experimental conditions (Tab. 1), the plasmid transfer rate is predominately controlled by the densities of D and R. Therefore, under such conditions, the excess in the availability of T has no impact on the estimated log*γ*. However, based on a preliminary survey using VME, we assume that at higher plasmid transfer rates (log*γ* > - 9) and in mediums that support high bacterial densities (Kc > 10E9 CFU/ml), *MAT* would have an increased impact on the estimated log*γ*. This is because, when the densities of all three strains tend to get large, the actual densities of mature T and D are important to avoid overestimation of T and underestimation of the log*γ*. Nevertheless, there is a need for further research to quantify the effect of *MAT* on the log*γ* under such extreme experimental conditions.

How general as these results likely to be? We expect our findings to be applicable for all typical well-mixed mating systems, which fall under the wide range of the experimental conditions (Tab. 1) covered in our study. Using the VME approach, we were able to quantify the effect of four common structural model deficits on the fitted plasmid transfer rate constants. The method has helped us to identify suitable sampling intervals for estimating the log*γ* in liquid culture mating experiments. Our study suggests that, under a wide range of experimental conditions, the lag phase and the plasmid cost are the most important additions to the original model by Levin et al. (1979). *LAG* and *CST* should be taken into account when estimating the rate constant of plasmid transfer from observations of D, R, and T in a well-mixed culture to limit the estimation error to an acceptable level (e.g., < 1 order of magnitude). In systems with short lag phases (< 0.5 hours), fast growth rates (*μ* > 2 hour^−1^), or an unbalanced initial D:R (D:R ≥ 100:1 or ≤ 1:100), *DNS* should not be neglected in a conjugation model. Based on the sensitivity analysis of the model outcome (estimated log*γ*) with respect to the various experimental conditions, we conclude that *MAT* can be excluded from the model without resulting in any significant errors in the parameter estimate. In such cases, T*young* + T*mature* should be modeled as a single variable - total T. Considering all the data of our study, sampling at a rate of 1 sample collected every 2 hours (for 12 hours), found to be sufficient for the proper inference of γ. This would also make sense in practice because too frequent sampling disturbs the culture repeatedly. While the effect of several structural deficits on estimates of plasmid transfer rate can also be predicted using logical reasoning, we need models that can: a) quantify the individual biases and their compensatory effect when used in combinations (for example, under default settings at sampling done every 2 hours, *LAG* as an individual process improved transfer rate estimate by 1.3 orders of magnitude, but when used in combination with *CST* and *DNS* improved the estimated log*γ* by 1 order, compare Fig. 2a with 2b) and b) analyze the sensitivity of the parameter estimates to different experimental conditions.

Our model accurately mimics plasmid dynamics in typical mating systems (Headd and Bradford, 2020; Kneis et al., 2019), but there are plasmids whose transfers are controlled in a different way and therefore result in processes that have an opposite impact on (i.e., overestimate) the plasmid transfer rate constant. For example, plasmid transfer can be induced by quorum sensing or pheromones (Koraimann and Wagner, 2014; Starčič et al., 2003), in the absence of nutrients (Headd and Bradford, 2018) or in the presence of antibiotics stress (Wistrand-Yuen et al., 2018). These processes that promote conjugation, when ignored from a model, would result in an overestimation of the plasmid transfer parameter. Similarly, transitory derepression, which causes a temporary increase in the transfer rate of the newly formed T by 2-6 orders of magnitude (Lundquist and Levin, 1986), would have an opposite impact compared to *MAT*. Simonsen et al. (1990) also studied the effect of excluding transitory depression from the transfer rates estimated through their analytical solution (end-point method) and observed an overestimation of the log*γ* by ≈ 1 order of magnitude in extreme situations (log*γ_derepressed_* = −9, duration of derepression = up to 6 generations, i.e., 2.5 hours for *μ* = 1.7 hours^-1^). We assume that under most of the real-world scenarios, the processes that we have ignored in our study would have a similar minimal impact on the improvement of the estimated plasmid transfer rate constant. This assumption is based on the weak dependence observed in this study between estimated plasmid transfer rate constants and the processes that affected conjugation rates only (e.g., *DNS* and *MAT*). Nevertheless, there is a need for further research to quantify the effect of the above-mentioned and other possible structural deficits on the estimates of plasmid transfer rate constants under different experimental conditions.

It is important to find the effect of various structural deficits on the model output to build IMs which generate non-biased parameter estimates. When an ill-structured inverse model is fitted to the experimental data, the estimated (reported) parameters might have large biases. These parameters would then generate bad predictions of strain densities when structurally accurate forward models are used for simulating the spread of ARG. On the other hand, if a wrongly fit model is used to estimate the parameters from empirical data, and then the same model is used for predictions of population densities, we might see a compensatory effect between several wrongly fit process rates, leading to correct prediction of some strains while others continue to be biased. For example, ignoring *CST* might lead to underestimation of the true log*γ*, but the T produced due to an increased growth rate and lower plasmid transfer rate might still be correctly estimated, whereas the D and R in this case with be grossly over and underestimated due to the wrongly fitted growth rate constant.

Virtual mating experiments can be used to compare and segregate the bias arising from known sources (e.g., data error and model structural error). The most important outcome of our study is the fact that, under all studied scenarios, the four structural deficits (*CST, LAG, DNS, MAT*) of the fitted models result in underestimation of the plasmid transfer rate constant. That negative bias in transfer rate constants may - at least in parts - explain why simulation models tend to predict a too quick disappearance of the plasmid-bearing strains in the absence of advantageous plasmid-encoded traits - a fact that is known as the “plasmid paradox” (Carroll and Wong, 2018). We also discussed other possible structural deficits which might have an opposite effect on the estimates of plasmid transfer rate constants. Because structurally deficient inverse models were used extensively in past decades, a considerable proportion of plasmid transfer rate constants reported in the literature is probably subject to biases. Our study demonstrates that the fitting of the structurally adequate inverse models can reduce the bias in estimated transfer rate constants and thus contribute to improved predictions of plasmid spread.

## Acknowledgments

The research leading to these results were supported by the German Federal Ministry of Education and Research [grant: 02WRS1377D (HyReKA)], the European Union [grant: 1822 (DSWAP)], and the TU Dresden Excellence initiative, which is funded by the German Federal and State Governments. We thank our colleague M. Zwanzig for the collaborative generation of some of the ideas developed in this study and U. Klümper, J. Feldbauer, and A. Chandrashekar for their valuable feedback during the development of this manuscript.

